# *In vitro* determination of the CB1 efficacy of illicit synthetic cannabinoids

**DOI:** 10.1101/385583

**Authors:** Shivani Sachdev, Kiran Vemuri, Samuel D. Banister, Mitchell Longworth, Michael Kassiou, Marina Santiago, Alexandros Makriyannis, Mark Connor

## Abstract

**BACKGROUND AND PURPOSE:** The morbidity and mortality associated with recreational use of synthetic cannabinoid receptor agonists (SCRAs) is a major health concern, and may involve over-activation of CB1 receptors. Thus, we sought to determine the efficacy of 13 SCRAs at CB1 using receptor depletion with the irreversible CB1 antagonist AM6544 followed by fitting the curve with the Black and Leff operational model to calculate efficacy.

**EXPERIMENTAL APPROACH:** Receptor depletion in mouse AtT-20 neuroblastoma cells stably expressing human CB1 was achieved by pre-treatment of cells with AM6544 (10 µM, 60 mins). The CB1-mediated hyperpolarisation of AtT20 cells was measured using membrane potential dye. From data fit to the operational model, the efficacy (*tau*) and affinity (K_A_) parameters were obtained for each drug.

**KEY RESULTS:** AM6544 did not affect the potency or maximal effect of native somatostatin receptor-induced hyperpolarisation (Control, pEC_50_ 9.13 ± 0.05, E_max_ 38 ± 1%; AM6544 treated pEC_50_ 9.18 ± 0.04, E_max_ 39 ± 0.7%). The *tau* value of ∆^9^-THC was 70-fold less than the reference CB-agonist CP55940, and 240-fold less than the highest efficacy SCRA, 5F-MDMB-PICA. Most of the SCRAs had about 50% of the efficacy of CP55940. There was no correlation between the *tau* and K_A_ values for any SCRA.

**CONCLUSION AND IMPLICATIONS:** All the SCRA tested showed substantially higher agonist activity at CB1 than ∆^9^-THC, which may contribute to the adverse effects seen with these drugs but not ∆^9^-THC, although the mechanisms underlying SCRA toxicity are still poorly defined.

## Introduction

Synthetic cannabinoid receptor agonists (SCRAs) are a large class of novel psychoactive substances (NPS), notionally designed to mimic the effects of tetrahydrocannabinol (∆^9^-THC), the main psychoactive ingredient in cannabis (Wiley et al., 2014). SCRAs have been marketed as herbal incense blends (often known as Spice or K2) and legal cannabis substitutes which are undetectable using conventional drug tests (Auwärter et al., 2009). Since the first generation of SCRAs (including JWH-018, JWH-073, JWH-200, and CP 47,497) were detected in herbal blends in 2008 (Auwärter et al., 2009, Teske et al., 2010), more than 230 SCRAs have been reported from 106 countries to the United Nations Office on Drugs and Crime (United Nations Office on Drugs and Crime, 2017). SCRA use has been associated with adverse health effects including hundreds of hospitalisations and dozens of fatalities (Cooper, 2016, Springer et al., 2016, Adams et al., 2017). The most commonly reported adverse effects are psychosis, anxiety, agitation, seizures, tachycardia, hypothermia, and kidney injury (Babi et al., 2017, Tait et al., 2016). In addition to these life-threatening effects, daily SCRA use has been linked to dependence and withdrawal (Macfarlane and Christie, 2015).

SCRAs activate cannabinoid type 1 (CB1) and type 2 (CB2) receptors, with their psychoactive effects derived from CB1 activation (Pacher et al., 2006). While cannabinoids have been reported to have activity at a variety of ion channels and GPCRs other than CB1 and CB2 (De Petrocellis and Di Marzo, 2010, Morales and Reggio, 2017), the relevance of these interactions for the effects of cannabinoids in humans remains to be established, and their potential role in SCRA toxicity is unknown. In rodents, both JWH-018- and AM-2201-induced seizures are mediated by CB1 (Malyshevskaya et al., 2017, Funada and Takebayashi-Ohsawa, 2018, Vigolo et al., 2015), while SCRA-induced hypothermia and bradycardia have also been shown to be CB1 dependent (Banister et al., 2016, Banister et al., 2015b, Banister et al., 2013). Intriguingly, a recent report suggests that the hypertensive effects of some SCRAs in rats may be independent of CB1 (Schindler et al., 2017).

While most SCRAs studied to date activate CB1 receptors with greater potency and efficacy than ∆^9^-THC in both [^35^S]GTPγS binding (Gamage et al., 2018, Wiley et al., 2015), and fluorescence-based membrane potential assays (Banister et al., 2013, Banister et al., 2015b), there is little quantitative information about the efficacy of SCRA at CB1. A substantial component of SCRA toxicity may be mediated through activation of CB1, and it is important to define the efficacy of SCRAs as a first step towards understanding possible mechanisms of CB1-mediated toxicity. The similar maximal effects of SCRAs reported in assays of membrane potential or [^35^S]GTPγS binding may be due to the presence of a receptor reserve, with only submaximal receptor occupancy by agonists needed to achieve their maximal response. Depleting receptor reserve can allow for quantitative determination of efficacy, by fitting concentration-response data before and after receptor depletion to the operational model of Black and Leff (Black and Leff, 1983). We have used the irreversible CB1 antagonist AM6544 (Finlay et al., 2017) that was able to deplete 94% of the receptor reserve available to effectively measure the efficacy of SCRAs to produce CB1-dependent hyperpolarisation of intact AtT-20 cells expressing human CB1. We determined the efficacy of a range of SCRAs identified in the NPS market since 2008 and found that the SCRAs we tested had up to 300 times the efficacy of ∆^9^-THC, with most having an efficacy about 50% of that of the reference agonist CP55940. While there was no obvious correlation between efficacy to hyperpolarise the cells and toxicity, we have established an assay that can be used to quantitate CB1 efficacy efficiently, and which is readily adaptable to the study of other CB1 signalling pathways.

## Methods

### Cell culture

Experiments utilised mouse AtT-20 pituitary tumour cells stably expressing human CB1 receptor as previously described (Banister et al., 2016). Cells were cultured in Dulbecco’s Modified Eagle Media (DMEM, Sigma-Aldrich, St. Louis, MO, USA) supplemented with 10% fetal bovine serum (FBS, Sigma-Aldrich, St. Louis, MO, USA), 100 unit/ml penicillin/streptomycin (Thermo Fischer Scientific, Waltham, MA, USA) and 80 µg/ml hygromycin (InvivoGen, San Diego, CA, USA). The cells were grown and maintained in 75cm^2^ flask and passaged at 80% confluency, or grown to 90% confluency for assay. Cells were incubated at 37 ^o^C in a humidified 5% CO_2_ atmosphere.

### Membrane potential assay

Changes in membrane potential of cells were measured using the fluorometric imaging plate reader (FLIPR) membrane potential (blue) assay kit (Molecular Devices, Sunnyvale, CA) as previously described (Knapman et al., 2013). Cells were detached from the flask using trypsin/EDTA (Sigma-Aldrich) and the pellet was resuspended in 10 ml Leibovitz’s (L-15) media supplemented with 1% FBS, 100 unit/ml penicillin/streptomycin and 15 mM glucose. The cells were seeded in a volume of 90 µl in poly D-lysine (Sigma-Aldrich) coated, black wall, clear bottom 96 well microplates. Cells were incubated overnight at 37 ^o^C in ambient CO_2_.

We initially tested the ability of the irreversible CB1 antagonist methyl arachidonyl fluorophosphonate (MAFP) to deplete the receptor reserve. On the following day, cells were incubated with MAFP (10 µM) for 20 mins at 37 ^o^C. MAFP was removed and the cells were washed twice with HBSS, which consisted of (mM) NaCl 145, HEPES 22, Na_2_HPO_4_ 0.338, NaHCO_3_ 4.17, KH_2_PO_4_ 0.441, MgSO_4_ 0.407, MgCl_2_ 0.493, CaCl_2_ 1.26, glucose 5.56 (pH 7.4, osmolarity 315 ± 5). Changes in membrane potential were measured after addition of CP55940 or somatotropin release-inhibiting factor (SRIF).

The novel CB1 irreversible antagonist AM6544 was synthesised at the Center for Discovery, Northeastern University, based on published procedures (Finlay et al., 2017). The day after plating, cells were incubated with AM6544 (10 µM) for 60 minutes, after removal of the L-15, at 37 ^o^C in ambient CO_2_. The cells were then washed twice with HBSS and loaded with 90 µl/well of L-15 media and 90 µl/well of reconstituted FLIPR dye. Control cells were treated the same way with HBSS alone. The AM6544-treated and control cells were compared side by side. The cell plate was incubated at 37 ^o^C in ambient CO_2_ for 1 hour prior to measuring the fluorescence using a FlexStation 3 microplate reader (Molecular Devices). The cells were excited at a wavelength of 530 nm and emission measured at 565 nm, with cut-off at 550 nm, and the readings were made every 2s. Baseline readings were taken for 2 mins after which 20 µl of drug was added to each well to give the desired concentration. The drugs of various concentrations were prepared in HBSS containing 0.1% BSA and 1% DMSO. The final concentration of DMSO in each well was always 0.1%. A concentration-response curve (CRC) for CP55940 was performed each day for quality control. On rare occasions, AM6544 pre-treatment failed to produce a substantial shift in CP55940 responses, probably indicating experimenter error, and these results were discarded.

SCRAs were synthesised as previously described in Banister et al. (2015a), (2015b, 2016). Chemical structure of SCRAs can be found in supplementary material (Supplementary Figure 1). We have previously shown that the effects of SCRAs in AtT20-CB1 cells were blocked by SR141716A, and that none of the SCRAs produced a significant change in the membrane potential of AtT20 wild-type cells (Banister et al., 2016, Banister et al., 2015b, Banister et al., 2015a, European Monitoring Centre for Drugs and Drug Addiction, 2018b). SCRA-mediated hyperpolarisation of AtT20-CB1 cells is also pertussis toxin (PTX) sensitive, confirming that the response is G_i/o_-dependent (Banister et al., 2016, Banister et al., 2015b, Banister et al., 2015a, European Monitoring Centre for Drugs and Drug Addiction, 2018b).

**Figure 1.**
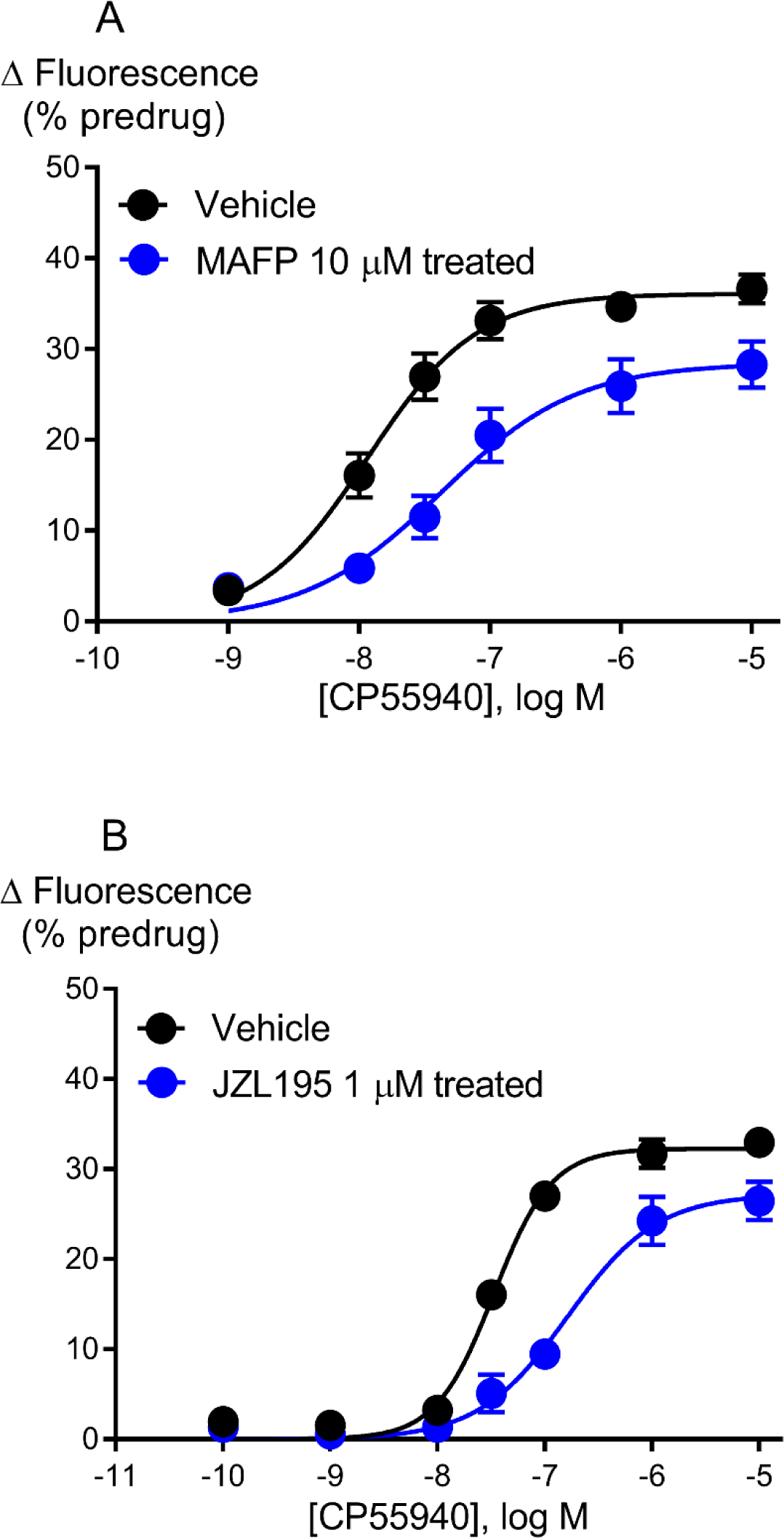
Concentration response curves for CP55940 mediated hyperpolarisation of AtT20-CB1 following pre-treatment with 10 µM of MAFP (A) and 1 µM of JZL195 (B). Representative data are presented as percentage in fluorescence corresponding to the hyperpolarisation of the cells. Each point represents the mean ± SEM of 6 independent determinants performed in duplicate and pooled data was fitted with four parametric logistic equation with bottom constrained to 0.

### Data Analysis

Drug responses are reported as percentage change of baseline fluorescence, following correction for the vehicle responses (0.1% DMSO). The hyperpolarisation of the cells produces a *decrease* in fluorescence. For convenience, values are expressed such that a change of 30% means a *reduction* in fluorescence of 30%. Data for individual experiments were analysed using the Black and Leff operational model in PRISM (Graph Pad Software Inc., San Diego, CA), using four-parameter non-linear regression to fit the operational model-receptor depletion equation. From the operational model, efficacy (*tau*) and affinity (K_A_) parameters were obtained for each drug. The basal parameter was constrained to zero, high efficacy agonists were included in each experiment, and their maximal response (E_max_) was used to define the E_max_ for the cells for that day. For individual drugs, K_A_ and n (transducer slope) parameters were shared between the AM6544-treated and control state (Equation 1). The *tau* parameter in the control state was used to measure the CB1 agonist efficacy. In this model of receptor activation, *tau* represents the inverse of the number of receptors that need to be occupied by an agonist to produce a response 50% of the system maximum. The transducer coefficient R [log(*tau*/K_a_)] was calculated for each ligand to include both efficacy and affinity simultaneously in a single parameter. Log(*tau*/K_a_) was estimated as log(*tau*) – log(K_A_).

The equation for operational model-depletion presented in the same style as Prism:

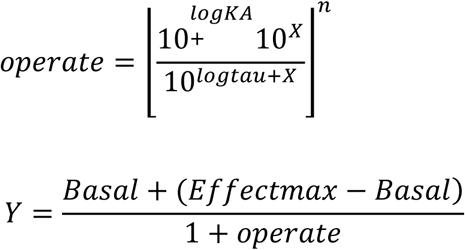

Unless otherwise noted, the data represent mean ± SEM of at least six independent experiments, each conducted in duplicate. The data and the statistical analysis comply with the recommendations on experimental design and analysis in pharmacology (Curtis et al., 2018, Curtis et al., 2015). Statistical significance is defined as P < 0.05.

### Nomenclature of targets and Ligands

Key protein targets and ligands in this article are hyperlinked to corresponding entries in http://www.guidetopharmacology.org/, the common portal of data from the IUPHAR/BPS Guide to PHARMACOLOGY (Harding et al., 2017), and are permanently archived in the Concise Guide to PHARMACOLOGY 2017/2018 (Alexander et al., 2017).

### Materials

CP55940, 2-arachidonolylglycerol, anandamide, MAFP, JZL195 and CUMYL-4CN-BINACA was purchased from Cayman Chemical Company (Ann Arbour, MI, USA); tetrahydrocannabinol was obtained from The Lambert Initiative (Sydney, NSW, Australia). AM6544 was a gift from laboratory of Professor Alexandros Makriyannis (Northeastern University, Massachusetts, USA). All the SCRAs unless otherwise stated, were synthesized by Samuel D. Banister and Mitchell Longworth in the lab of Michael Kassiou at Sydney University (Sydney, NSW, Australia). All the drugs were stored in aliquots of 30 mM at −80 °C until needed.

## Results

### MAFP as a CB1 irreversible antagonist

We initially examined whether MAFP, which has been reported to irreversibly antagonise native CB1 receptor function (Fernando and Pertwee, 1997), was an appropriate antagonist for the present study. Pre-treatment with MAFP (10 µM, 20 min) resulted in a reduction in the hyperpolarisation induced by subsequent application of maximally effective concentration of CP55940 (10 µM) compared to untreated cells (Figure 1A, Control E_max_ 36 ± 1.3% and MAFP treated E_max_ 29 ± 2.2%), consistent with reduction in receptor number. MAFP pre-treatment did not significantly affect the hyperpolarisation induced by SRIF (100 nM) in the same cells (Supplementary Figure 2, P > 0.05), suggesting that it did not interfere with G-protein coupled receptor signalling to G-protein gated inward rectifying potassium (GIRK) channels *per se*. However, MAFP inhibits a range of enzymes, including those that degrade endocannabinoids such as, fatty acid amide hydrolase (FAAH) and monacylglycerol lipase (MAGL) (Goparaju et al., 1999). To assess whether the decrease in response of CP55940 may have been related to changes in endocannabinoid levels, we investigated the effects of the structurally unrelated, non-selective inhibitor of FAAH and MAGL, JZL195 (Long et al., 2009), on CP55940 signalling. Pre-treatment of cells with JZL195 (1 µM, 60 min) also inhibited the hyperpolarisation induced by CP55940 (Figure 1B, Control E_max_ 32 ± 0.8% and JZL treated E_max_ 27 ± 1.8%), suggesting that altering endocannabinoid degradation can alter CB1 signalling; therefore, due to uncertainty surrounding the mechanism through which MAFP alters CP55940 responses, we turned to AM6544 to deplete the CB1 receptors.

**Figure 2.**
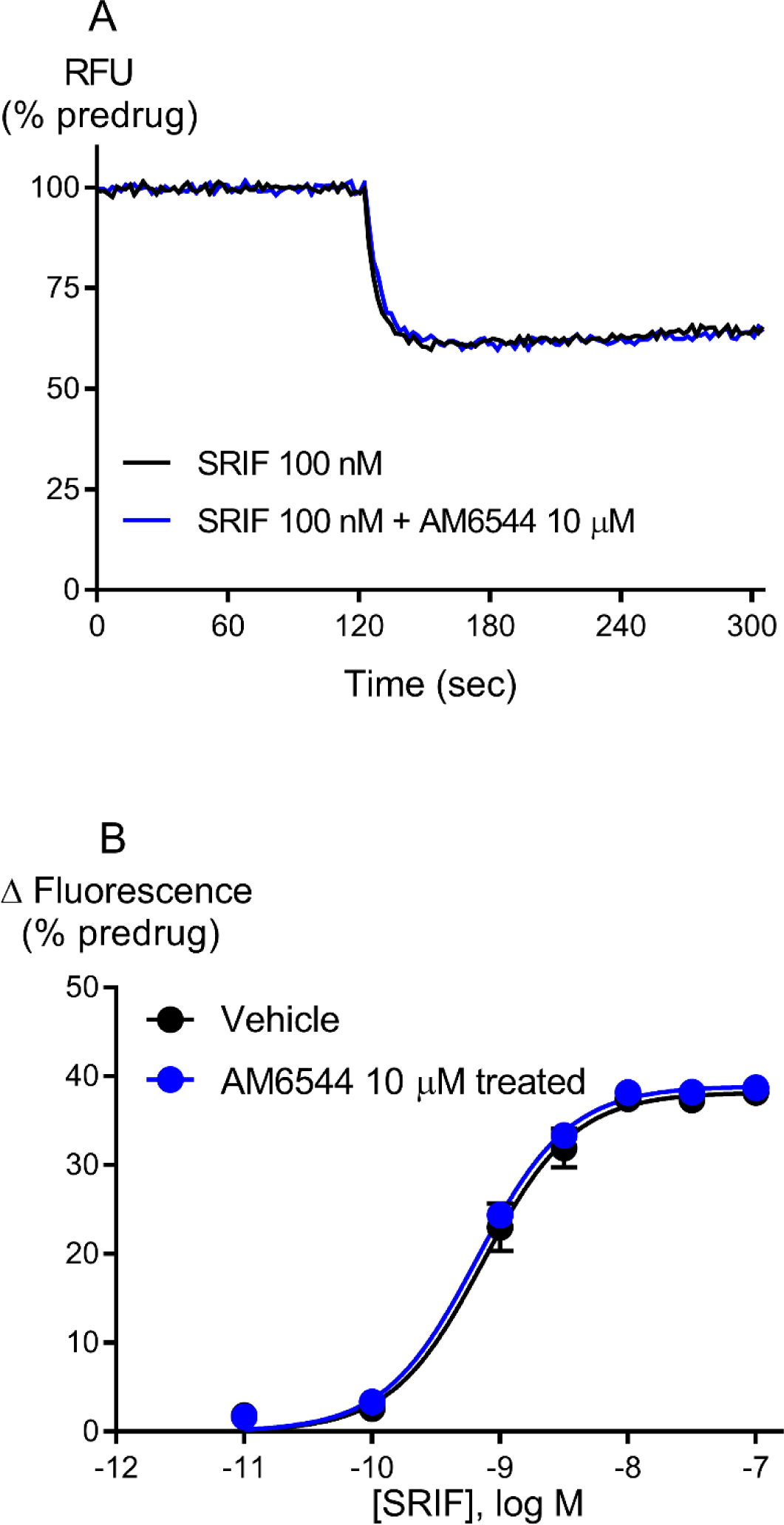
AM6544 is a specific irreversible antagonist of CB1 receptor. (A) Raw trace showing the change in fluorescence normalised to the predrug baseline for SRIF on AtT20-CB1 cells pre-treated for 60 min with vehicle or AM6544 (10 µM) and then washed twice before incubation with MPA dye. The traces are representative of at least six independent experiments. (B) Concentration response curve for SRIF mediated hyperpolarisation of AtT20-CB1 following pre-treatment with AM6544 (10 µM) or vehicle. Data represents the mean ± SEM of 6 independent determinants performed in duplicate. There was no difference in the potency or maximal effect of SRIF between vehicle or following pre-treatment with AM6544.

### Specificity of AM6544: a novel CB1 irreversible antagonist

*In vitro*, AM6544 behaves as an irreversible antagonist of CB1 (Finlay et al., 2017). To confirm that AM6544 does not non-specifically interfere with receptor signalling mechanisms in AtT20-CB1, we examined the effect of AM6544 on the activation of native SRIF receptors. Pre-treatment with AM6544 (10 µM, 60 min) had no effect on the potency or maximal effect of SRIF induced hyperpolarisation when compared to vehicle treated cells (Control, pEC_50_ 9.13 ± 0.05, E_max_ 38 ± 1%; AM6544 treated pEC_50_ 9.18 ± 0.04, E_max_ 39 ± 0.7%, Figure 2), indicating that AM6544 did not interfere with either SRIF receptors or their signalling pathways, likely shared with CB1.

### Functional activity of cannabinoids after receptor depletion with AM6544

The efficacy of the classical CB1 agonist, CP55940, was measured after the pharmacological knockdown of CB1 receptors with AM6544 (10 µM, 60 min). The maximal response of CP55940 (10 µM) was reduced after AM6544 pre-treatment compared to vehicle treated cells (Control, E_max_ 33 ± 2; AM6544 treated E_max_ 26 ± 2, P < 0.05, Figure 3A). The *tau* value for CP55940 was reduced 14-fold in AM6544 pre-treated cells compared to vehicle cells (Control, *tau* 91 ± 21; AM6544 treated *tau* 6 ± 2, Table 1), suggesting that AM6544 can effectively deplete the receptors available to high efficacy SCRAs. From the operational model, the pK_A_ of CP55940 was estimated to be 5.4 ± 0.1 (Table 1, n = 17).

**Table 1.** Efficacy and Functional affinity of CP55940, ∆^9^-THC and other SCRAs. Values were calculated using the operational model of pharmacological agonism following CB1 receptor depletion with AM6544, as outlined in the Methods.

| Compound | Operational efficacy tau (±SEM) | Functional affinity pK<sub>A</sub> (±SEM) | Log (tau/K<sub>A</sub>) | Transducer slope (n) |
| --- | --- | --- | --- | --- |
| CP55940 | 91 (22) | 5.37 (0.12) | 7.53 (0.06) | 0.91 (0.07) |
| WIN55212-2 | 24 (19) | 5.27 (0.35) | 6.79 (0.08) | 1.77 (0.29) |
| $\Delta^9$-THC | 1.3 (0.3) | 5.65 (0.30) | 6.18 (0.21) | 0.83 (0.17) |
| 2-AG | 36 (19) | 4.98 (0.13) | 6.61 (0.20) | 0.92 (0.08) |
| AEA | 3 (0.5) | 5.12 (0.14) | 5.69 (0.08) | 1.06 (0.13) |
| JWH-018 | 25 (8) | 5.91 (0.17) | 7.39 (0.09) | 0.98 (0.78) |
| AM-2201 | 18 (3) | 6.74 (0.10) | 8.06 (0.08) | 1.12 (0.11) |
| UR-144 | 22 (8) | 4.94 (0.17) | 6.31 (0.11) | 1.28 (0.09) |
| XLR-11 | 34 (10) | 5.61 (0.19) | 7.32 (0.14) | 1.33 (0.04) |
| PB-22 | 60 (28) | 6.58 (0.16) | 8.34 (0.04) | 1.04 (0.15) |
| 5F-PB-22 | 54 (15) | 7.22 (0.10) | 8.95 (0.09) | 1.16 (0.08) |
| AB-CHMINACA | 42 (23) | 7.28 (0.09) | 8.69 (0.12) | 1.29 (0.12) |
| AB-PINACA | 34 (19) | 6.48 (0.30) | 8.51 (0.13) | 1.10 (0.12) |
| MDMB-CHMICA | 82 (10) | 6.36 (0.04) | 8.26 (0.06) | 1.14 (0.16) |
| MDMB-FUBINACA | 309 (168) | 6.72 (0.20) | 8.98 (0.11) | 0.91 (0.10) |
| 5F-MDMB-PICA | 314 (155) | 6.55 (0.24) | 9.12 (0.06) | 1.23 (0.09) |
| CUMYL-4CN-BINACA | 45 (15) | 7.61 (0.11) | 9.24 (0.09) | 1.18 (0.14) |

**Figure 3.**
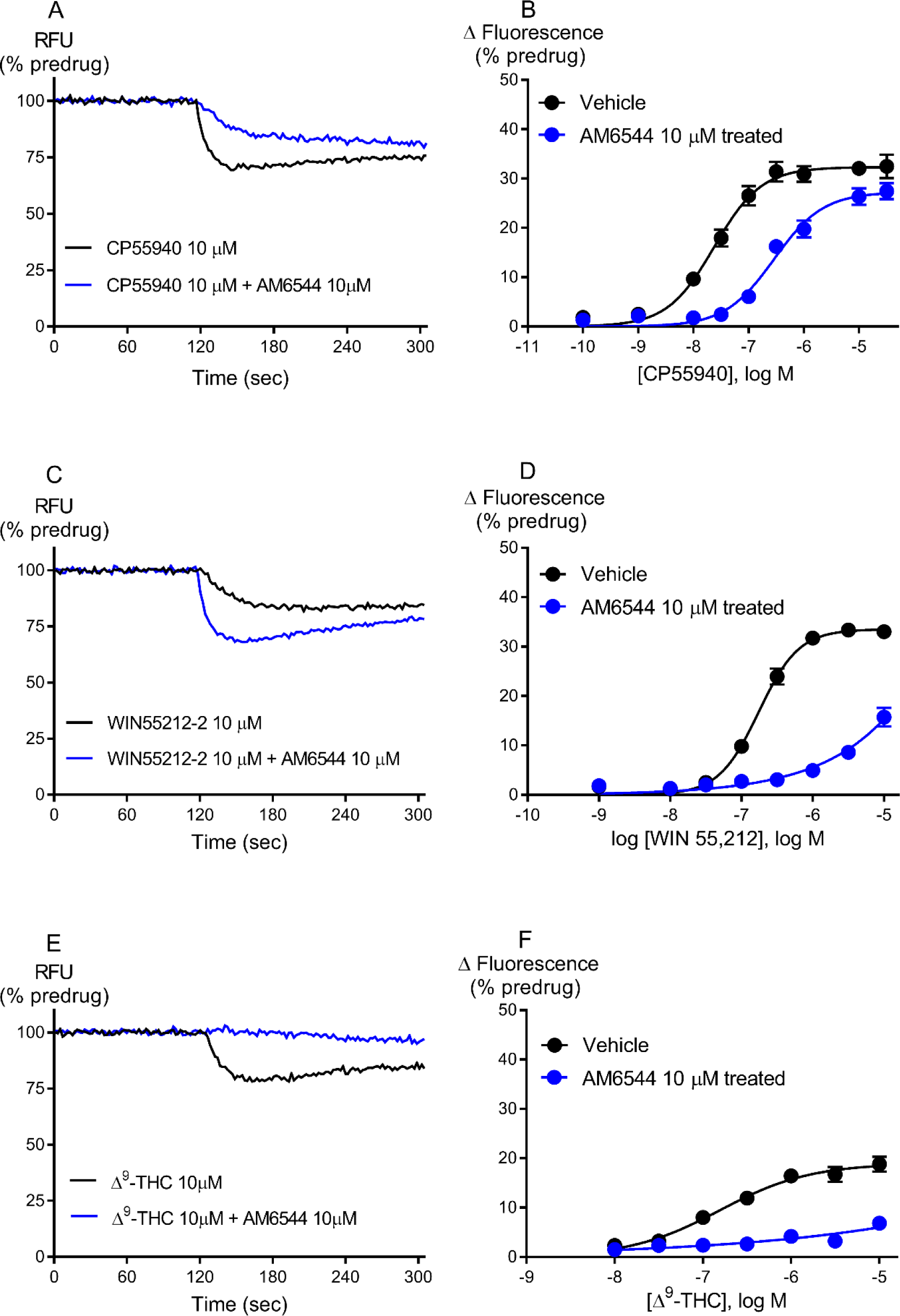
Representative traces for reference compounds CP55940 (A), WIN55212-2 (C) and ∆^9^-THC (E) following pre-treatment with vehicle or AM6544 (10 µM) on AtT20-CB1 cells. Raw trace showing reduction in hyperpolarisation induced by maximally effective concentration (10 µM) of CP55940, WIN55212-2 and ∆^9^-THC after AM6544 pre-treatment compared to vehicle. Concentration response curves for CP55940 (B), WIN55212-2 (D) and ∆^9^-THC (F) were plotted using four parameter nonlinear regression to fit the operational model-receptor depletion equation with basal constrained to 0. Data represents the mean ± SEM of at least 6 independent determinants performed in duplicate.

We also determined the efficacy of WIN55212-2, ∆^9^-THC and endogenous cannabinoids on CB1 after receptor depletion with AM6544. The hyperpolarisation produced by WIN55212-2 was strongly inhibited by AM6544 pre-treatment (10 µM, 60 min) compared to vehicle treated cells (Figure 3). The *tau* for WIN55212-2 was reduced 4-fold compared to CP55940, but was 19-fold greater than ∆^9^-THC (Table 1). The efficacy of endogenous cannabinoids, 2-arachidonolylglycerol and anandamide were respectively 3- and 28-fold less than that of CP55940 (Figure 4, Table 1).

**Figure 4.**
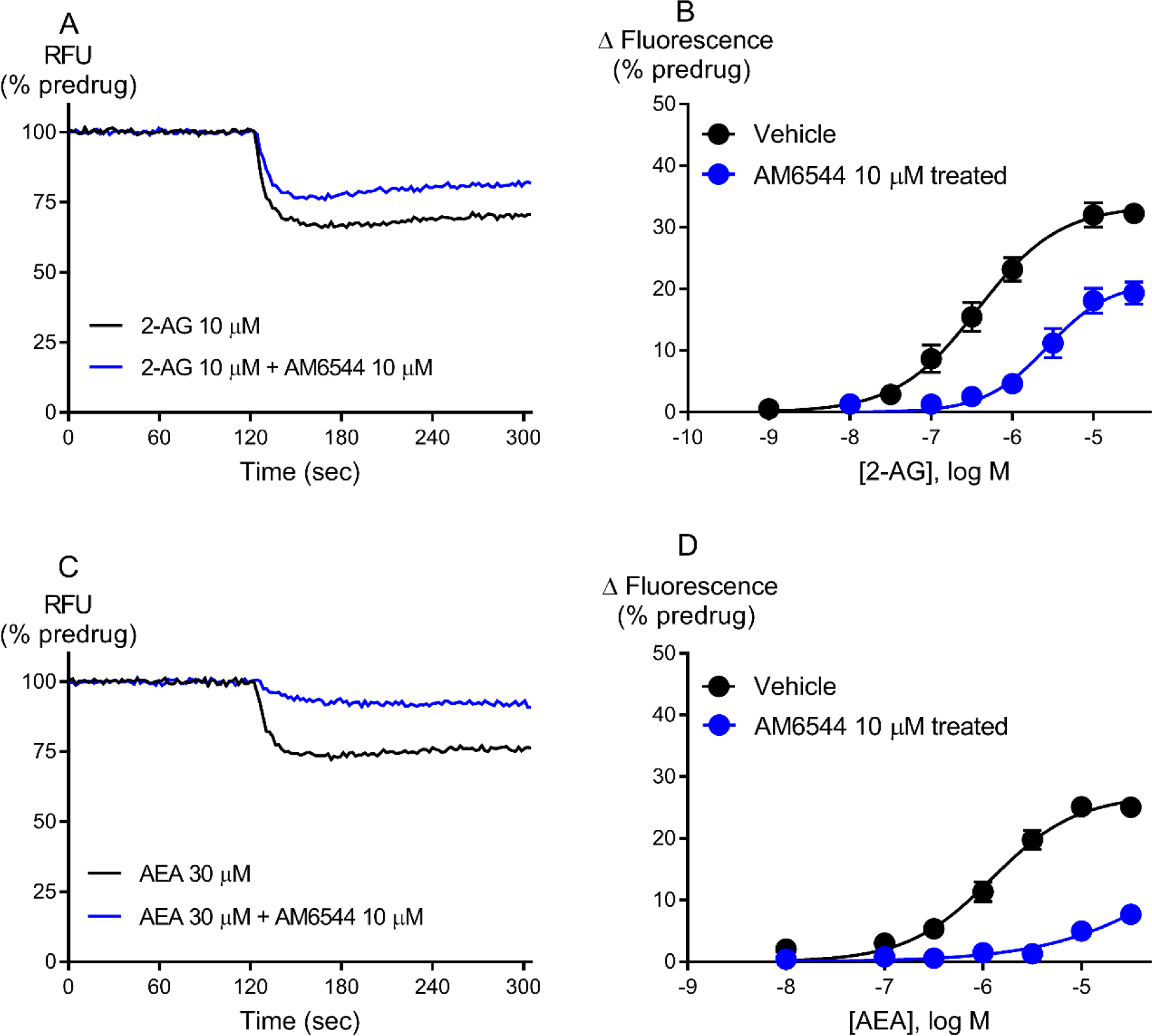
Representative traces for endogenous cannabinoids 2-arachidonolylglycerol, 2-AG (A) and anandamide, AEA (C) after pre-treatment with vehicle or AM6544 (10 µM) on AtT20-CB1 cells. Raw trace showing reduction in hyperpolarisation induced by maximally effective concentration of 2-AG (10 µM) and AEA (30 µM) after AM6544 pre-treatment compared to vehicle. Concentration response curves for 2-AG (B) and AEA (D) were plotted using four parameter nonlinear regression to fit the operational model-receptor depletion equation with basal constrained to 0. Illustrates the increase in efficacy of 2-AG as compared to AEA. Data represents the mean ± SEM of at least 6 independent determinants performed in duplicate.

We assessed the relative efficacy of SCRAs to provide novel insight into potential mechanisms of toxicity and the evolution of SCRAs structures over time. The efficacy of SCRAs was determined following receptor depletion with AM6544. Example traces and CRC are shown for JWH-018, MDMB-FUBINACA and CUMYL-4CN-BINACA (Figure 5). The efficacy for all the drugs we examined are found in Table 1. The *tau* of SCRAs tested ranged from 18 to 314 with two of the 13 SCRAs having *tau* values greater than 300 (5F-MDMB-PICA and MDMB-FUBINACA). The first SCRA to be identified in Spice (Atwood et al., 2010, Banister and Connor, 2018), JWH-018, exhibited 3.7-fold less *tau* than CP55940 but 18-fold higher *tau* than ∆^9^-THC. Most of the SCRAs tested, including the most recently identified compound available for this study (CUMYL-4CN-BINACA) had approximately 50% of the efficacy of CP55940 (Table 1). The least efficacious SCRA, AM-2201, showed 5-fold less *tau* than CP55940 while the most efficacious SCRA, 5F-MDMB-PICA, showed 4-fold higher *tau* than CP55940 (Table 1).

**Figure 5.**
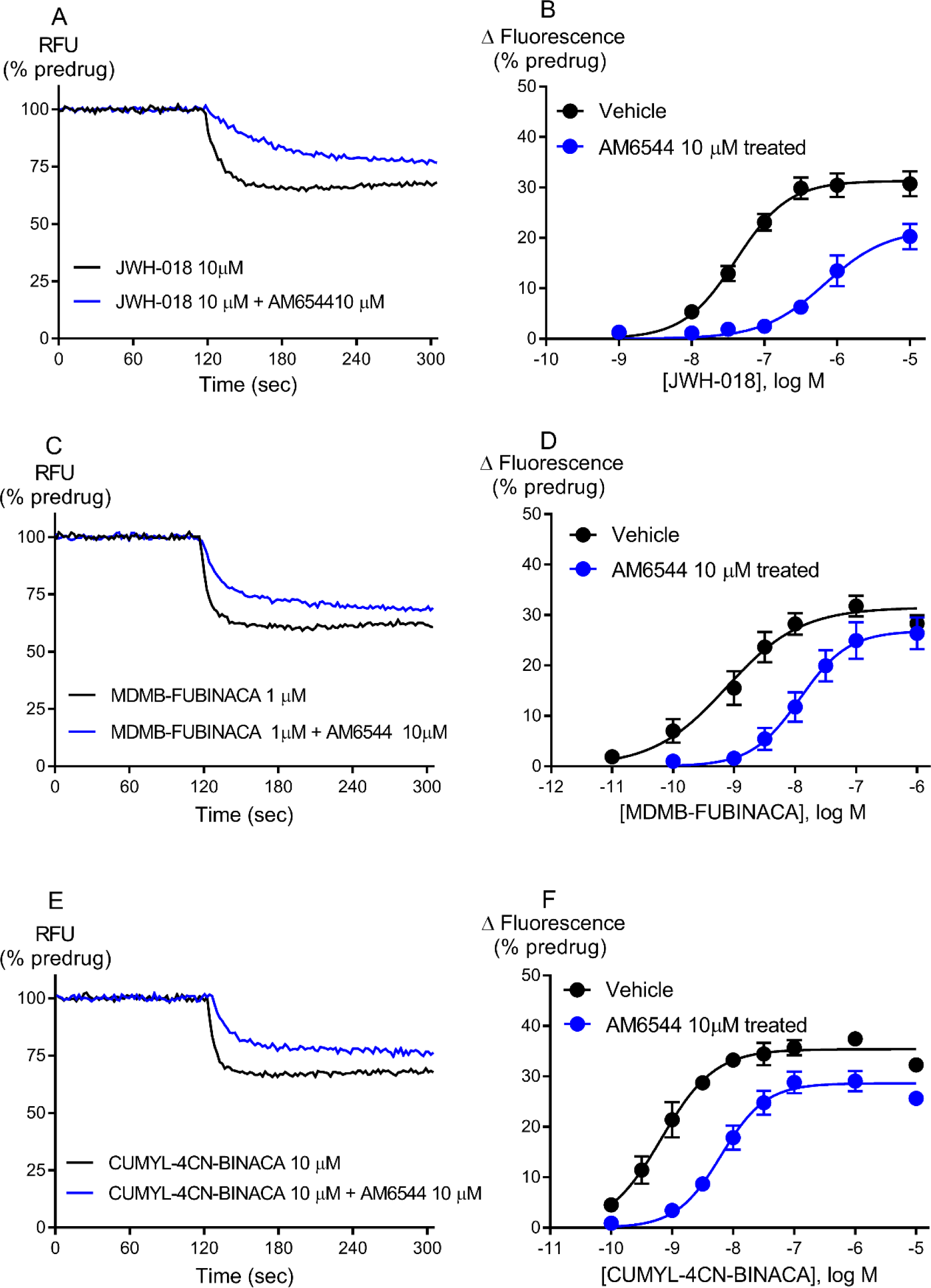
Representative traces for JWH-018 (A), MDMB-FUBINACA (C) and CUMYL-4CN-BINACA (E) following pre-treatment with vehicle or AM6544 (10 µM) on AtT20-CB1 cells. Raw trace showing reduction in hyperpolarisation induced by maximally effective concentration of JWH-018 (10 µM), MDMB-FUBINACA (1 µM) and CUMYL-4CN-BINACA (10 µM) after AM6544 pre-treatment compared to vehicle. Concentration response curves for JWH-018 (B), MDMB-FUBINACA (D) and CUMYL-4CN-BINACA (F) were plotted using four parameter nonlinear regression to fit the operational model-receptor depletion equation with basal constrained to 0. Data represents the mean ± SEM of at least 6 independent determinants performed in duplicate.

We calculated the functional affinity of SCRAs at CB1 using the operational model (Table 1). In the present study, the K_A_ of SCRAs ranged from 25 nM to 12 µM, where three of the 13 SCRAs had K_A_ values less than 100 nM (CUMYL-4CN-BINACA, AB-CHMINACA and 5F-PB-22) and three demonstrated micromolar affinities (JWH-018, XLR-11 and UR-144). Most of the SCRA had a higher affinity for CB1 compared to CP55940, with the exception of WIN55212-2 and UR-144, which had 1.5- and 3-fold lower affinity respectively (Table 1). No correlation was found between the operational efficacy and affinity obtained for SCRAs (Figure 6, r^2^ = 0.06, P > 0.05).

**Figure 6.**
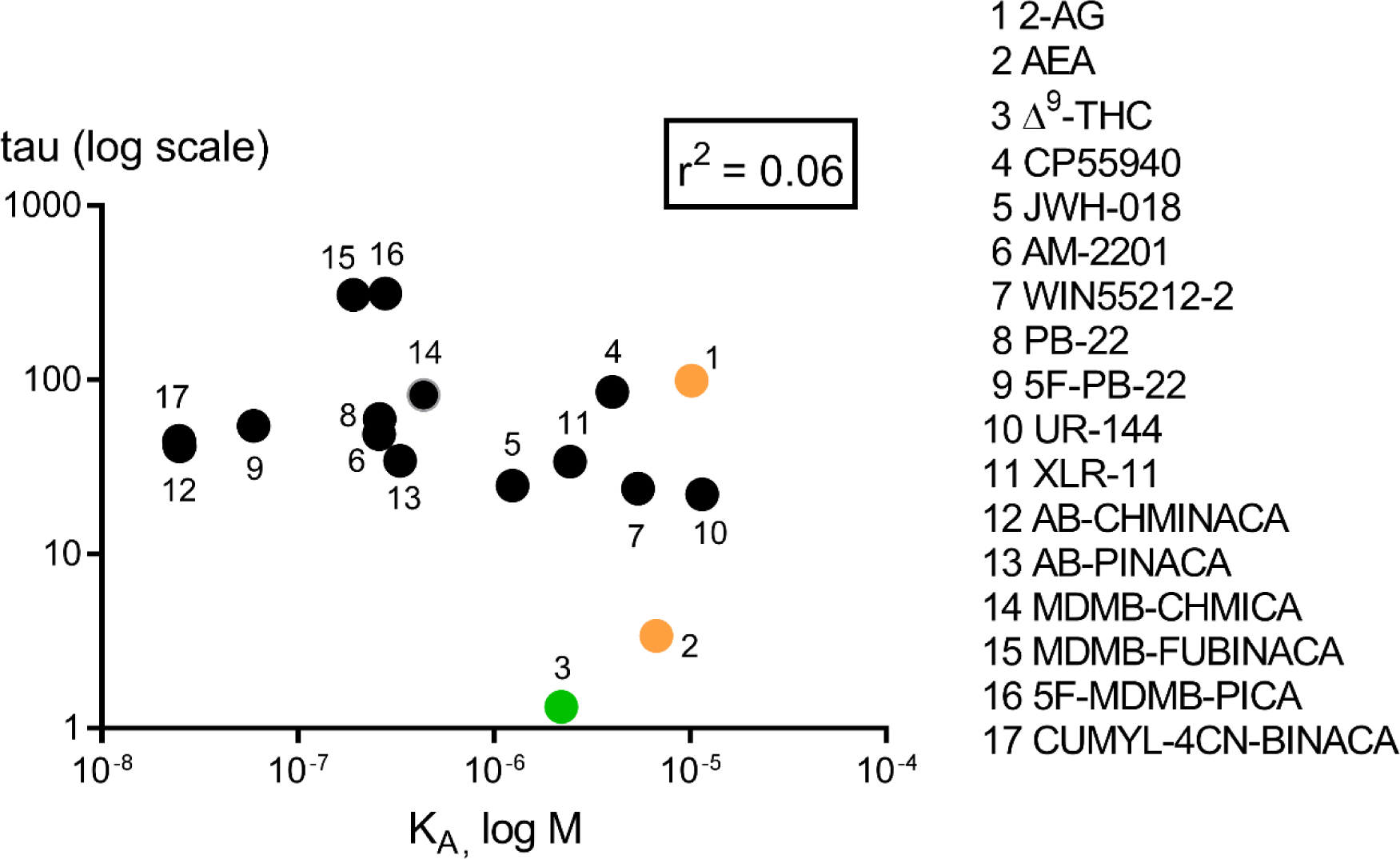
Correlation of operational efficacy (tau) and functional affinity (K_A_) for CB1 agonists on Gi dependent activation of GIRK channel in AtT20 cells. Representative data are presented, demonstrating an r^2^ of 0.06 (p > 0.05), where tau and K_A_ values shown are the fitted values from the operational analysis.

We compared the maximal activity of SCRAs on G_i/o_ pathway via Gβγ-dependent activation of GIRK channels, by calculating the log (*tau*/K_A_) for each of the SCRAs as described in the methods. Two of the SCRAs (5F-MDMB-PICA and MDMB-FUBINACA) exhibited 4-fold higher operational efficacy (log tau) than most of the SCRAs tested. However, when data was measured using log (*tau*/K_A_) scale, four of the 13 SCRAs had log (*tau*/K_A_) values more than 8.9 (CUMYL-4CN-BINACA, 5F-MDMB-PICA, MDMB-FUBINACA, and 5F-PB-22), suggesting that the overall maximal intrinsic effect is produced by both efficacy and affinity.

To determine the percentage of CB1 receptors available after AM6544 pre-treatment, the ratio of *tau* post- and pre-receptor depletion were measured for each SCRA. The average value of *tau* post- and pre-receptor depletion curves for SCRAs tested was found to be 0.06 ± 0.05, indicating that AM6544 caused an overall 94% reduction in receptors available to CB1 agonists. To further understand the extent by which AM6544 is depleting CB1, the *tau* value of 5F-MDMB-PICA (highest efficacy) was compared to ∆^9^-THC (lowest efficacy). The *tau* value for 5F-MDMB-PICA was roughly 242-fold higher than ∆^9^-THC, suggesting that the *tau* of higher efficacy SCRAs can only be quantified with substantial depletion of receptor reserve.

## Discussion

This study has measured the efficacy of a wide range of SCRAs in a naturalistic signalling assay in intact cells. Following receptor depletion with the irreversible CB1 antagonist AM6544, we were able to use the Black and Leff operational model to calculate the efficacy (*tau*) and affinity (K_A_) of these SCRAs. All the SCRAs tested showed substantially higher agonist activity at CB1 than ∆^9^-THC (*tau*, 1.3 ± 0.3), with *tau* that ranged between 18 and 314. Most of the SCRAs tested (JWH-018, AM-2201, WIN55212-2, PB-22, 5F-PB-22, UR-144, XLR-11, AB-CHMINACA, AB-PINACA, and CUMYL-4CN-BINACA) had approximately 50% of the efficacy of the widely used, non-selective cannabinoid CP55940. 5F-MDMB-PICA and MDMB-FUBINACA exhibited the highest efficacies from the SCRAs tested; however, there was no correlation between the *tau* and K_A_ of SCRAs or there is no obvious trend for decreasing/increasing *tau* over time. No obvious correlation was observed between SCRA efficacy to activate native GIRK channels and their reported adverse effects.

We have used the new CB1 irreversible antagonist, AM6544, that specifically depletes the CB1 receptor reserve from the pool available for orthosteric agonist binding. The specificity of AM6544 was confirmed by assessing the effect of AM6544 pre-treatment on the activation of native SRIF receptors in the same cells. We found that AM6544 did not interfere with SRIF receptors or their shared signalling pathways with CB1 reinforcing that AM6544 is a reliable pharmacological tool to antagonise the CB1-mediated effects.

The efficacy of SCRAs have principally been measured using [^35^S]GTPγS binding assays, which measure the accumulated activation of G proteins in membranes over a period of 30-60 minutes (Wiley et al., 2015, Gamage et al., 2018, Thomas et al., 2017). In these assays, the maximum response is used as the measure of efficacy, with the assumption that this maximum response is not constrained - that there is an excess of G-proteins relative to CB1 receptors. Given the high levels of receptor expression that can be achieved in recombinant systems, this assumption may not be valid (Gamage et al., 2018). In our study we have circumvented this limitation by reducing receptor number, and we have been able to measure a very wide range of apparent efficacies to produce acute hyperpolarisation of AtT-20-CB1 cells (> 300-fold), compared with a 2-3-fold difference in the maximum response to agonists in CB1 GTPγS assays (Table 2). For all SCRAs except MDMB-FUBINACA, the *E*_max_ in the GTPγS experiments is higher, relative to that of CP55940, than the *tau* value in the hyperpolarisation assay. While our assay seems sensitive to differences in efficacy, it represents activation of one pathway, likely by Gβγ-subunit activation of GIRK (Mackie et al., 1995). GTPγS assays provide a more general measure of Gα_i/o_-subunit activation, but uncoupled from signalling pathways (Ibsen et al., 2017). Although, there is intriguing evidence in the mouse for WIN55212-2 induced activation of GIRK channel in a CB1 dependent manner (Marinelli et al., 2009), these have not yet been demonstrated in human neurons. Neither assay effectively captures CB1 coupling to Gα_s_ or Gα_q_, or non-G protein mediated pathways, such as those dependent on arrestin. A more useful and transferable measure of intrinsic efficacy could be to use the transduction coefficient [log(*tau*/K_A_)] (Kenakin et al., 2012). An analysis of the log(*tau*/K_A_) values of the SCRAs we tested indicates that CUMYL-4CN-BINACA, 5F-MDMB-PICA, MDMB-FUBINACA, and 5F-PB-22 have an overall higher intrinsic efficacy on the Gα_i/o_ pathway compared to the other SCRAs tested. This may provide a more accurate reflection of the functional activity of the SCRAs to the activation of CB1-dependent native G-protein gated channel.

**Table 2.** Comparison of human CB1 functional efficacy for selected SCRAs at CB1 measured using [^35^S]GTPγS binding assay and FLIPR membrane potential assay The maximum response of SCRAs are expressed relative to CP55940 is shown in bold below the SEM For each measure the number of replicates *n*, is shown below the relative efficacy NA, not applicable MPA, membrane potential assay a values from (Gamage et al., 2018) represent E_max_ (95% confidence interval) for percentage increase over basal stimulation b values from (Ford et al., 2017) represent E_max_ (±SEM) are presented as the fraction of the effect produced by reference agonist CP55,940. c values from (Thomas et al., 2017) represent E_max_ (95% confidence interval) for percentage [^35^S]GTPγS with basal globally shared at 100% d values from (Wiley et al., 2015) represent E_max_ (±SEM) for percentage increase over basal stimulation

| Compound | [<sup>35</sup>S]GTPγS binding, E<sub>max</sub> | MPA, Operational efficacy tau (±SEM) |
| --- | --- | --- |
| Δ<sup>9</sup>-THC | 32 (27-36)<sup>a</sup><br>**0.53**<br>3 | 1.3 (0.3)<br>**0.01**<br>6 |
| JWH-018 | 1.02 (± 0.10)<sup>b</sup><br>**NA**<br>4 | 25 (± 8)<br>**0.27**<br>6 |
| UR-144 | 193 (164-221)<sup>c</sup><br>**0.91**<br>5 | 22 (± 8)<br>**0.24**<br>6 |
| XLR-11 | 205 (177-233)<sup>c</sup><br>**0.97**<br>5 | 34 (± 9.9)<br>**0.37**<br>6 |
| PB-22 | 415 (373-458)<sup>c</sup><br>**1.97**<br>2 | 60 (± 28)<br>**0.66**<br>6 |
| AB-CHMINACA | 205 (± 14)<sup>d</sup><br>**1.65**<br>6 | 42 (± 23)<br>**0.46**<br>6 |
| AB-PINACA | 192 (± 25)<sup>d</sup><br>**1.55**<br>6 | 34 (± 19)<br>**0.38**<br>6 |
| MDMB-FUBINACA | 75 (68-82)<sup>a</sup><br>**1.25**<br>3 | 309 (± 168)<br>**3.41**<br>6 |

NA, not applicable
MPA, membrane potential assay

The rapid emergence of SCRAs as recreational drugs is a major public health concern as little is known about their pharmacology and their toxicological profiles are inconsistent and unpredictable (Castaneto et al., 2014, Zawilska and Andrzejczak, 2015, Frinculescu et al., 2017). The use of SCRAs is associated with a wide variety of adverse effects in humans (Sherpa et al., 2015, Adams et al., 2017). These effects may be mediated via both CB1 and non-CB1 systems (Kaushik et al., 2011, Sherpa et al., 2015, Silva et al., 2018). All 13 SCRAs tested in this study had a much higher efficacy than ∆^9^-THC, suggesting that adverse effects produced by ∆^9^-THC intake may not provide a good guide to the potential consequences of SCRA high efficacy CB1 activation. SCRAs produce CB1-mediated seizures in animals, in addition to the well characterised CB1-mediated “tetrad” of hypolocomotion, catalepsy, anti-nociception, and hypothermia, and could conceivably account for some of the adverse effects of SCRAs in humans (Vigolo et al., 2015). We also measured the efficacy of two principal endocannabinoids, 2-AG and AEA. Our results are consistent with previous reports, showing that 2-AG is a higher efficacy agonist of CB1 compared to AEA (Di Marzo and De Petrocellis, 2012), and that both have a higher efficacy than ∆^9^-THC (Pertwee, 1997). Thus ∆^9^-THC, but not most of the SCRAs investigated here, is likely to act as an antagonist of 2-AG modulation of neuronal activity *in vivo* (Straiker and Mackie, 2005, Pertwee, 2008).

Cannabinoid interactions with renal and cardiovascular systems have also been described (Pacher et al., 2005), but the degree to which these interactions are influenced by agonist efficacy is unknown. A specific toxicity attributed to a particular SCRA was the acute kidney injury linked to the use of XLR-11 (Thornton et al., 2013). The present study shows that XLR-11 exhibited about 10-fold less efficacy than the highest efficacy SCRA tested, 5F-MDMB-PICA, which has not been reported to produce kidney injury. XLR-11 affects kidney cells via CB receptors on mitochondria, rather than through the plasma-membrane delimited pathway we have examined, and both CB1 and CB2 receptors were reported to be involved in the toxic effects of XLR-11 *in vitro* (Silva et al., 2018). Thus, toxicity for individual SCRAs potentially involves a complex interplay between activity at both CB1 and CB2 receptors, efficacy at CB1 receptors, cellular and subcellular distribution, and access to receptors to different body and cellular compartments, and the formation of bioactive drug metabolites (Fantegrossi et al., 2014).

The emergence of NPS provides a continual challenge to the development of targeted interventions and novel therapeutics to help minimise the adverse effects associated with their use (European Monitoring Centre for Drugs and Drug Addiction, 2018a). Although some SCRAs were mined from older patents (AM2201, AB-CHMINACA, AB-FUBINACA, UR144, etc.), newer drugs have unprecedented structures (Banister and Connor, 2018). We assessed a diversity of SCRAs from the earliest to most recent examples identified in the NPS market. There was no obvious trend for decreasing/increasing *tau* over time and *tau* or functional affinity, suggesting that SCRAs are not designed to be more efficacious over time. Our data also show no correlation between SCRAs efficacy and functional affinity for CB1. While it is not immediately apparent what causes the toxic effects of SCRAs and whether signalling of SCRAs at Gα_s_, Gα_q_ or arrestins rather than Gα_i/o_ dependent CB1 signalling is important. However, it is clear that these drugs are likely to have very different pharmacological profiles to the commonly consumed cannabinoid, ∆^9^-THC. This was demonstrated in a recent study where AB-CHMINACA showed specific CB1-dependent activation of Gα_s_ signalling (Costain et al., 2018). These observations highlight the complexity of the pharmacology of SCRAs mediated activation of different signalling pathways downstream of CB1. Thus, more work is needed to better understand the physiological consequences of signalling pathways resulting from CB1 activation by high efficacy agonists.

## Acknowledgements

This work was supported by NHMRC Project Grant 1107088 awarded to M.K., and M.C. and National Institutes of Health grant P01DA009158 to A.M. Work was also supported in part by the European Union’s Seventh Framework Programme [FP7/2007-103] InMind (Grant Agreement No. HEALTH-F2-2011-278850). S.S. was supported by a Macquarie University Doctoral Scholarship. I would like to thank Dr. Chris Bladen and Alexander Gillis for their input in reading and commenting on draft work.

## Author contributions

S.S. designed and performed experiments, analysed the data and wrote the manuscript. Data analysis was performed by M.C. and S.S. V.K.V. and A.M. generated the AM6544 compound. The synthesis of drugs was carried out by S.D.B. and M.L. with direction from M.K. M.S. made and characterised CB1 cells. The manuscript was drafted by S.S. and M.C. with contributions from S.D.B., V.K.V., and M.K. M.C. supervised the study and revised the manuscript. All the authors have given approval to the final version of manuscript.

## Abbreviations

SCRA: synthetic cannabinoid receptor agonists
HA: haemagglutinin
CB1: cannabinoid receptor type 1
AtT20-CB1: mouse pituitary tumour cells stably transfected with HA tagged human CB1 receptors
NPS: novel psychoactive substances
GIRK: G protein-coupled inwardly-rectifying potassium channel
SRIF: somatotropin release-inhibiting factor
MAFP: methyl arachidonyl fluorophosphonate
PTX: pertussis toxin
FAAH: fatty acid amide hydrolase
MAGL: monacylglycerol lipase

